# An efficient method of evaluating multiple concurrent management actions on invasive populations

**DOI:** 10.1101/2021.07.28.452079

**Authors:** Amy J. Davis, Randy Farrar, Brad Jump, Parker Hall, Travis Guerrant, Kim M. Pepin

**Affiliations:** National Wildlife Research Center, Wildlife Services, Animal and Plant Health Inspection Service, United States Department of Agriculture, 4101 Laporte Ave., Fort Collins, CO, 80521, USA; Wildlife Services, Animal Plant Health Inspection Service, United States Department of Agriculture, Puxico, MO; Wildlife Services, Animal Plant Health Inspection Service, United States Department of Agriculture, Marshfield, MO, 65706; Wildlife Services, Animal Plant Health Inspection Service, United States Department of Agriculture, Gainesville, FL, 32641; Wildlife Services, Animal Plant Health Inspection Service, United States Department of Agriculture, Columbia, MO, 65202

## Abstract

Evaluation of the efficacy of management actions to control invasive species is crucial for maintaining funding and to provide feedback for the continual improvement of management efficiency. However, it is often difficult to assess the efficacy of control methods due to limited resources for monitoring. Many managers view effort on monitoring as effort taken away from performing management actions. We developed a method to estimate invasive species abundance, evaluate management effectiveness, and evaluate population growth overtime from a combination of removal activities (e.g., aerial gunning, trapping, ground shooting) using only data collected during removal efforts (the method of removal, the date, location, number of animals removed, and the effort). This dynamic approach allows for estimating abundance at discrete time points and the estimation of population growth between removal periods. We applied this method to removal data from Mingo National Wildlife Refuge in Missouri from December 2015 to September 2019, where the management objective is elimination. Populations of feral swine on Mingo NWR have fluctuated over time but showed more marked declines in the last 3-6 months of the time series. More dramatic declines were observed in the center of the refuge likely aided by reducing potential immigration from the rest of the refuge. To counteract population growth (from both births and immigration) the percent of the population of feral swine removed monthly must be greater than the growth rate. On average, we found that removing 10% of the population monthly had only a 50% chance of causing a population decline, whereas removing 15% of the population monthly had a 95% chance of causing a population decline. Our method provides advancement over traditional removal modeling approaches because it can be applied to evaluate management programs that use a broad range of removal techniques concurrently and whose management effort and spatial coverage vary across time.

## INTRODUCTION

Invasive species pose significant threats to native ecosystems, human health, and the global economy (Pejchar and Mooney 2009, Early et al. 2016, Paini et al. 2016). Efficient methods for control of invasive species are critical to mediate the increasing challenges they present. However, it is challenging to assess the efficacy of control methods because of the trade-off in effort aimed at performing management actions and effort aimed at collecting monitoring data to evaluate management; decisions to divert resources from removal efforts to other activities such as monitoring may be met with opposition. Yet, regular evaluation provides evidence of the impact that resources spent on control activities have on reducing invasive species and feedback for the continual improvement of efficient management actions.

Population evaluations for wildlife are often conducted using methods that require monitoring data in addition to records of management actions (mark-recapture, transect sampling, etc.). However, ideally it is possible to evaluate the effects of management actions on population abundance using records of removal efforts and removal models (Zippin 1958). The benefit of this approach is that resources would not be diverted from critical control activities while information for improving management outcomes could still be gleaned. Removal models only require simple information which is routinely collected during management (i.e. number of animals removed, the effort used to remove the animals, location, and date/time) and are commonly used for population evaluation in pest or harvested species of fish and wildlife (Zippin 1958, Pollock 1991, Williams et al. 2002), including invasive species (Davis et al. 2016).

Management of invasive species often involves a suite of management techniques to take advantage of the fact that some methods perform better in a given habitat than others, or at different times of year, or involve different personnel or resource needs. Typically, removal models only consider removal data from a single removal source to adhere to the assumption of a constant capture rate. Therefore, modifications are needed to the standard removal model to be more broadly applicable to the range of methods used in invasive species management, and range of environments where these methods are applied. Drawing from multi-methods analyses (i.e., analyses of monitoring data that are derived from multiple sampling techniques; e.g., Nichols et al. 2008), it is possible to estimate detection rates (or capture rates) separately for each monitoring or capture method while using these distinct sources of data to inform the overall biological state of the system (Davis et al. 2019a). Applying multi-method analyses to removal models poses some unique challenges in that multiple removal events are needed to have the same probability of impacting all individuals in the population across the removal types. Given that some removal methods (e.g., trapping, ground shooting, aerial gunning) are unlikely to impact all individuals in the population similarly due to differences in the area impacted by a given removal device, adjustments need to be made on how to incorporate multiple removal methods.

Removal models work on the premise that – assuming a closed population and constant capture rate – the ratio of the number of animals removed in two subsequent removal events will reflect the ratio of the populations available to be removed at those two removal events. Removal models have been applied to populations of fish (Riley and Fausch 1992, Rosenberger and Dunham 2005), birds (Farnsworth et al. 2002, Alldredge et al. 2007), mammals (Andrea et al. 2007, Sullivan and Sullivan 2013), and particularly wild pigs (Waithman et al. 1999, Parkes et al. 2010, Davis et al. 2019b). Classic removal models are static and estimate abundance only at one point in time. However, most management for invasive species is conducted over many months or years and is conducted continuously. Therefore, a logical advancement to removal models is to integrate removal data into a robust design approach that allows for abundance estimation during a period of demographic closure, allowing for population growth between closed periods (Link et al. 2018), and estimation of population growth across time. Knowing the amount of population growth is important to ensure removal levels are sufficient to cause populations to decline and not simply keep pace with population growth (i.e., maintain a constant abundance).

Our goal was to develop a method to estimate abundance of invasive species, and to determine management intensities necessary for achieving management objectives, using only management data (i.e., without a separate monitoring effort). In addition to relying solely on removal data, we wanted to create a model that would: 1) incorporate all removal methods employed by managers, 2) be able to compare the efficacy of different removal methods (i.e. evaluate the capture rates of different removal methods), 3) evaluate population growth and compare it to removal levels, and 4) provide evaluation of management actions (i.e. determine if management efforts are sufficient to address management objectives).

## METHODS

### Study areas

Mingo National Wildlife Refuge (NWR) is located in south eastern Missouri. The refuge comprises of roughly 87 km^2^ of bottomland hardwood forest, cypress-tupelo swamp, marsh, and upland forest ecosystems.

### Data

For our study, we used data on feral swine (*Sus scrofa*) removal efforts to demonstrate the efficacy of this analytical approach. Feral swine are invasive in North America and are actively managed to reduce human-wildlife conflict and damage to agriculture, natural resources, and personal property. As part of the management program at Mingo NWR we recorded: the date of management actions, the location (latitude/longitude), the type of removal event (e.g., aerial gunning, trapping, ground shooting), the number of animals removed, and the effort involved (e.g., number of hours in a helicopter, number of trap nights, number of hours ground shooting). We used feral swine removal data from December 2015 to September 2019 from Mingo NWR. The removal efforts conducted on Mingo NWR included aerial gunning, trapping, and ground shooting.

We estimated capture rates for each method separately. These rates were allowed to vary depending on the effort applied by a method at any point in time (similar to St. Clair et al. 2012, Davis et al. 2016). For example, during aerial gunning events the number of hours per flight are recorded and thus an hourly capture rate was estimated. For trapping, there were multiple nights when more than one trap was active. Therefore, we used trap nights (number of active traps on a given night) as the base capture rate for trapping. Ground shooting effort was not easy to calculate since some ground shooting events are opportunistic and some are intentional. As capture numbers are generally low for ground shooting and because ground shooting events are often undertaken to find and remove a particular individual, hours spent searching is not a good measure of effort. Instead, we estimated the ground shooting capture rate as the number of independent ground shooting events that occur in a single day.

### Analytical methods

We subdivided our study areas into management units (sites) which were no larger than 150 km^2^ including a 2 km buffer (to account for the average maximum distance moved in a day for feral swine; Kay et al. 2017). We developed a dynamic removal model that incorporates multiple removal methods using a Bayesian hierarchical framework (Fig. 1). In keeping with removal model assumptions (Zippin 1958), we assumed the abundance for a primary study period, which we defined as one month (*t*), within a site (*i*) was closed to demographic changes, but allowed demography to vary month to month (i.e., assuming an open population over the full time frame of the study). Similar to St. Clair et al. (2012), we estimated site level abundance (*n*) and capture rate (*p*) from removal data using a multinomial distribution (Equation 1). This format models the number of animals removed (*y*_*ijtk*_) at site *i*, removal pass *j*, at time *t*, and from removal method *k*, as a function of the total number of animals in the population (*n*_*it*_) at site *i* and time *t*, and the probability of encounter (*π*_*jitk*_). The probability of encounter as described by St. Clair et al. (2012), is the probability that an animal is removed during pass *j* but not before that period. One difficulty with including multiple removal approaches in a single removal modeling framework, is that the area impacted by different removal methods is not the same (e.g., the area covered by aerial gunning compared to the area covered by a single trap). This disconnect would violate the assumption that all animals are equally catchable in the study area. Therefore, we applied the concept of availability (Diefenbach et al. 2007) to address this problem. The availability concept assumes that a portion of the population may not be available for detection by a surveillance approach. We equated the proportion of the area impacted by a given method to the proportion of the population that was available to be captured by that method (Equation 2; e.g., Pavlacky et al. 2012). If availability is not accounted for population estimates will be biased low. Therefore, we revised the probability of encounter to be the probability that an animal was removed and available on pass *j* but was either not available or available and not captured prior to pass *j* within time *t* (Equation 3; *l* in the equation represents values earlier than the current *j*). We estimated capture rate per unit effort per area for removal method *k* (*p*_*k*_) and estimated the capture rate accounting for effort per area (*g*_*ijk*_/*A*_*ijtk*_) as the cumulative capture rate (θ_ijtk_) for site *i*, removal pass *j*, time *t*, and method *k* (Equation 4). The area impacted was different for each removal method. In our study there were three removal methods: aerial gunning, trapping, and ground shooting. Aerial gunning was assumed to cover the entire study area. For traps, we used a buffer area around the active traps based on the area of influence of traps for feral swine calculated by McRae et al. (2020). For ground shooting a standard 5 km^2^ area was used per ground shooting event.

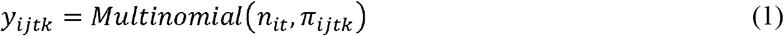

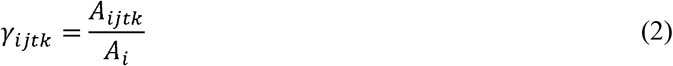

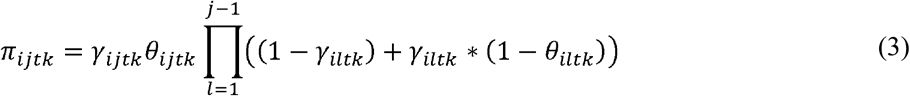

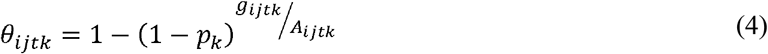

**Figure 1.**
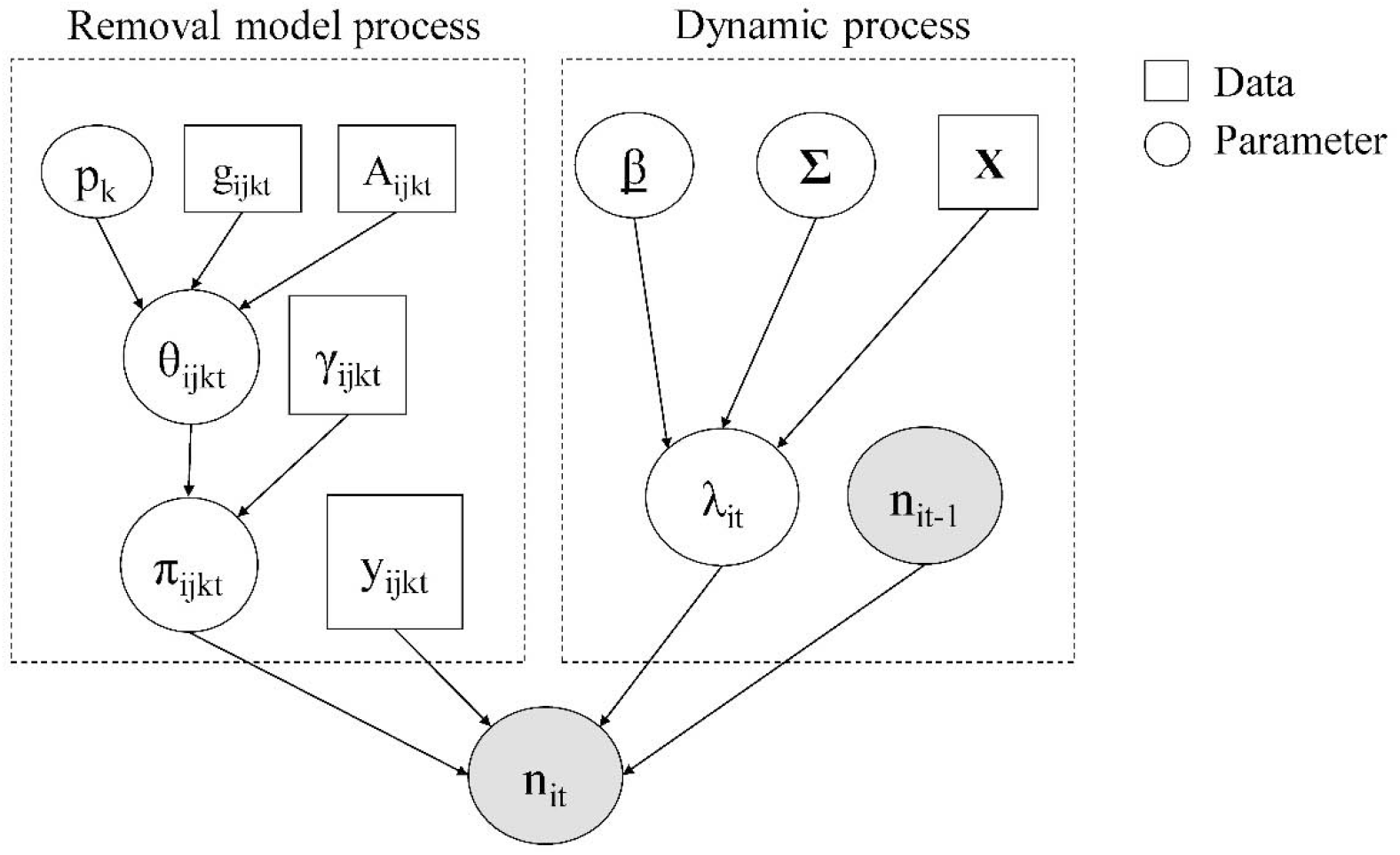
Directed acyclic graph (DAG) of the dynamic removal model with multiple removal methods that we applied to feral swine management data from Mingo National Wildlife Refuge. The squares represent data inputs, the circles represent parameters that are estimated in the model. The subscripts denote the site (*i*), the removal pass (*j*), the removal method (*k*), and the month (*t*). Abundance, *n*_*it*_, at each month (*t*) is estimated using an removal model where the number of animals removed, *y*_*ikjt*_, at site (*i*), removal pass (*j*), removal method (*k*), in month (*t*) is a function of the conditional capture rate (equation 3). This is a Malthusian growth model where the abundance, *n*_*it*_, at time *t* is dependent on the abundance at time *t−1*, based on the monthly growth rate (lambda it). Therefore, we include n_it−1_ as an input to n_it_, but show both abundance estimates (*n*) as shaded to denote the iterative nature of this parameter.

We modeled the logit of capture rates by method (*p*_*k*_) using a normal distribution with mean (*μ*_*p*_) and variance (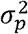; Equation 5). To model abundance across time we used an exponential growth model where the abundance remaining after removals at time *t−1* is multiplied by the growth rate at site *i* and time *t* (λ_*it*_; Equation 6). We modeled the abundance (*n*_*it*_) for site *i* and time *t* using a Poisson distribution (Equation 6). We modeled the growth rate (λ_*it*_) as a log-normal distribution (i.e. exponential population growth) with the mean representing a linear combination of covariates (***X***; Equation 7) with coefficients (**β**) modeled as a standard normal distribution (Equation 8). We allowed growth rate to vary across time using basis functions (Hefley et al. 2017).

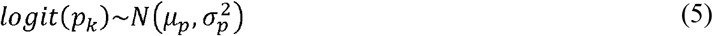

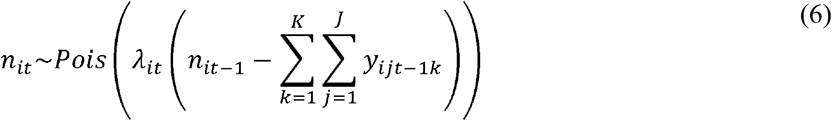

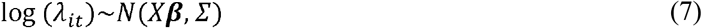

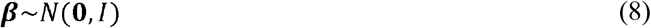

We used R (R Core Team 2017) to custom code a Markov chain Monte Carlo (MCMC) to fit the joint posterior distribution of the parameters of interest (Appendix A). We used several Metropolis-Hastings steps to model conditional distributions that were not identifiable distributions. Convergence was assessed graphically by visually assessing the trace plots for mixing and convergence. Posterior estimates are based on 50,000 MCMC iterations with the first 20,000 as burn-in.

Using estimates of the capture rate and the proportion of the population that needs to be removed per month, we can calculate the effort needed by each removal method to achieve the observed population reduction. To calculate the effort needed by each method we need to account for the estimated capture rate and the proportion of the total study area that an individual removal event per method would impact. The entire area covered by all study areas in Mingo NWR was 311.6 km^2^. Therefore, the proportion of the entire area impacted by each method was 5% for one trap, 13% for one hour of aerial gunning, and 2% for one ground shooting event. By setting the effective capture rate (*θ*_*k*_) to the mean instantaneous rate of increase (*r*_*it*_ = *log(λ*_*it*_)) we used Equation 4 to calculate for the effort needed per method (*g*_*k*_) to combat that growth.

### Simulation Analysis

We assessed the model’s performance by simulating true abundance and removal data and evaluating how well removal-based estimates of abundance recaptured the simulated (‘true’) abundance values. The objective of the simulation is to determine the conditions under which the model performs well and when it performs poorly, to provide guidelines on when this approach could be used with confidence. We simulated single method approaches to determine the impact of specific capture rates and percentages of area available on the reliability of the estimates. We examined the impact of capture rate, percent of area available, the number of months in the study, the number of removal passes by month, and the total area of the study on model performance. We ran all combinations of model parameters with 5 iterations each. We compared model performance using three metrics: 1) determining if the true abundance was within the 95% credible interval of the estimated abundance, 2) determining if the estimated abundance was within 10% of true abundance, and 3) examining the correlation between the estimates and truth across time. The first metric is a simple binary metric that determines if our estimate captures truth. However, this approach may over- or under-estimate the performance of the model if the error around the estimates are particularly narrow or wide. Therefore, we also wanted to see if the estimate was consistently within 10% of the truth, showing how well we match truth without relying on the amount of uncertainty in the abundance estimates. Our third metric looks at how well we match the overall trend the in the population. In some situations, the model may over or under-estimate the abundance routinely, but it may follow patterns in abundance well. These three metrics of success give information of the conditions under which the model performs well and when it does not.

Using these three metrics of success we identify conditions under which the model performs well. We were interested in criteria that we could determine from the input data only so the suitability of this approach could be determined from the data before conducting the analysis. For example, the input data would include the number of removals, the type of removal, the duration of the study, and the area of the study, they would not include the capture rate or the abundance which are estimated in the model. As removal models use a pattern of declining captures in their calculations, we calculated the trend in removal passes within a month to see if this pattern would indicate the data are suitable to use this approach.

## RESULTS

### Simulation Results

We simulated over one million months of removal data under a variety of conditions. The goal was to determine the range of conditions under which this method performed well, and therefore, we examined more conditions that may result in poorer data than conditions that would result in better data.

Almost half of the simulations resulted in true abundances that were less than 30 individuals. Removal model are known to perform poorly when estimating small abundances. Therefore, we focused on the simulations where the true abundance was above 30 individuals. Of the simulations with a true abundance of 30 or greater, 81% where within 10% of truth, 8% were biased high, and 11% were biased low. Only 41% of the true abundances were within the 95% credible intervals. However, 91% of simulations had a 90% or greater correlation with truth. This suggests that under many conditions the population estimate may be biased either high or low, but under most conditions this approach consistently does a good job of tracking trends in abundance across time. Estimates were more likely to be unbiased for studies that were a year or more in length. 61% of simulations that were 3 months long resulted in unbiased estimates compared to 86% for year long studies, and 93% for two year studies. Five or more removal events within a month performed better than when only three removal events were conducted. The size of the study area did not greatly impact how well the estimates performed, however, very low sizes (30 km^2^) and very large sizes (300 km^2^) were slightly worse. A larger impact on the reliability of the estimate was what proportion of the study area was impacted by the removal methods, in particular, if the removal methods only impacted 30% or less of the study area the estimates were more likely to be biased. The estimates also performed better as the capture rate increased. Capture rate is a combination of the ability of a method to capture animals (per unit capture rate) and the number of capture devices or amount of time spent by method (effort). Therefore, even when using a capture method that has a low individual capture rate (ground shooting for example), the cumulative capture rate can be increased by increasing effort.

Correlation between estimates and truth was high for the majority of simulated conditions. The only conditions where the correlation was less than 0.9 were when all of these conditions were met: having fewer than 21 animals removed within a study area in a month, having removal methods cover less than 30% of the study area, having a large study area (150 km^2^ or larger) and being only a six month or shorter study. These conditions would suggest that the number of animals removed relative to the abundance is so small that many combinations of abundance and capture rates could fit those data and thus the results are not reliable.

### Case study results

We estimated the population abundance of feral swine in three regions of Mingo NWR from December 2015 to September 2019. The total number of feral swine removed across the entire refuge during the study period was 2,926. The number removed was relatively consistent per year from 2016-2018 (523 in 2016, 488 in 2017, and 685 in 2018). 2019 saw the most removals at 1,213. Population estimates show relatively stable abundance for the majority of the study period. Only subpopulations 1 and 2 show declines and they were mostly within the last six months to a year of the study (Fig. 2).

**Figure 2.**
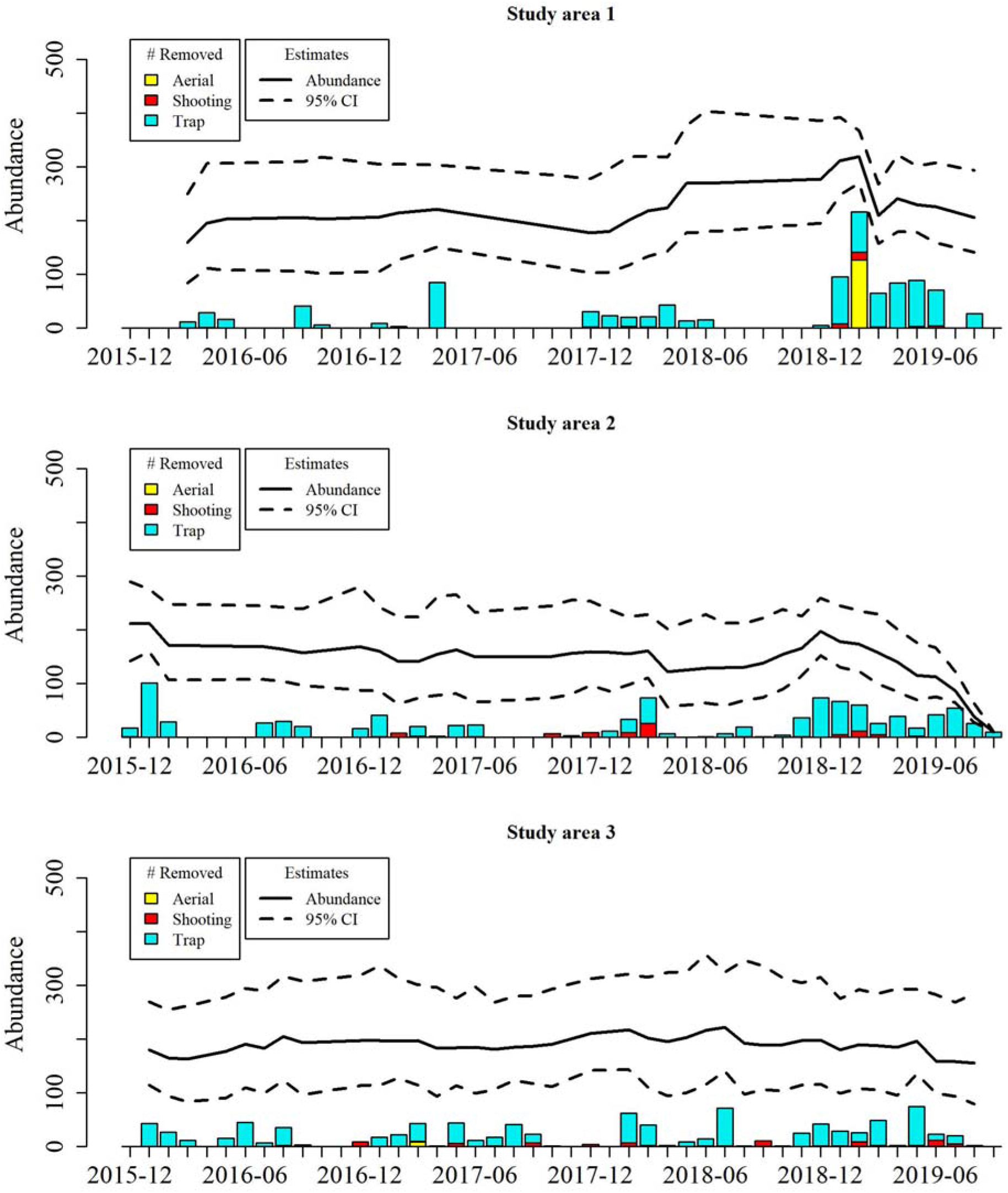
Feral swine abundance estimates (solid black line) and 95% confidence interval (dashed line) for three study areas of Mingo National Wildlife Refuge. The number of animals removed by month are shown as bars, colors indicate the method of removal (yellow for aerial gunning, red for ground shooting, and cyan for trapping).

In our case study, we found that on average the proportion of the population removed each month was 0.12 (95% CI: 0.08, 0.18) but varied by study area and year (Fig. 3). However, monthly growth from births and immigration was estimated at 1.17 on average, suggesting that for the first year the removals were working to hold the abundance constant rather than pushing the populations towards elimination (Fig. 3). The increased removal effort in the last six months of the study period was sufficient to combat the population growth and correspondingly cause population declines in two of the subpopulations (Fig. 3).

**Figure 3.**
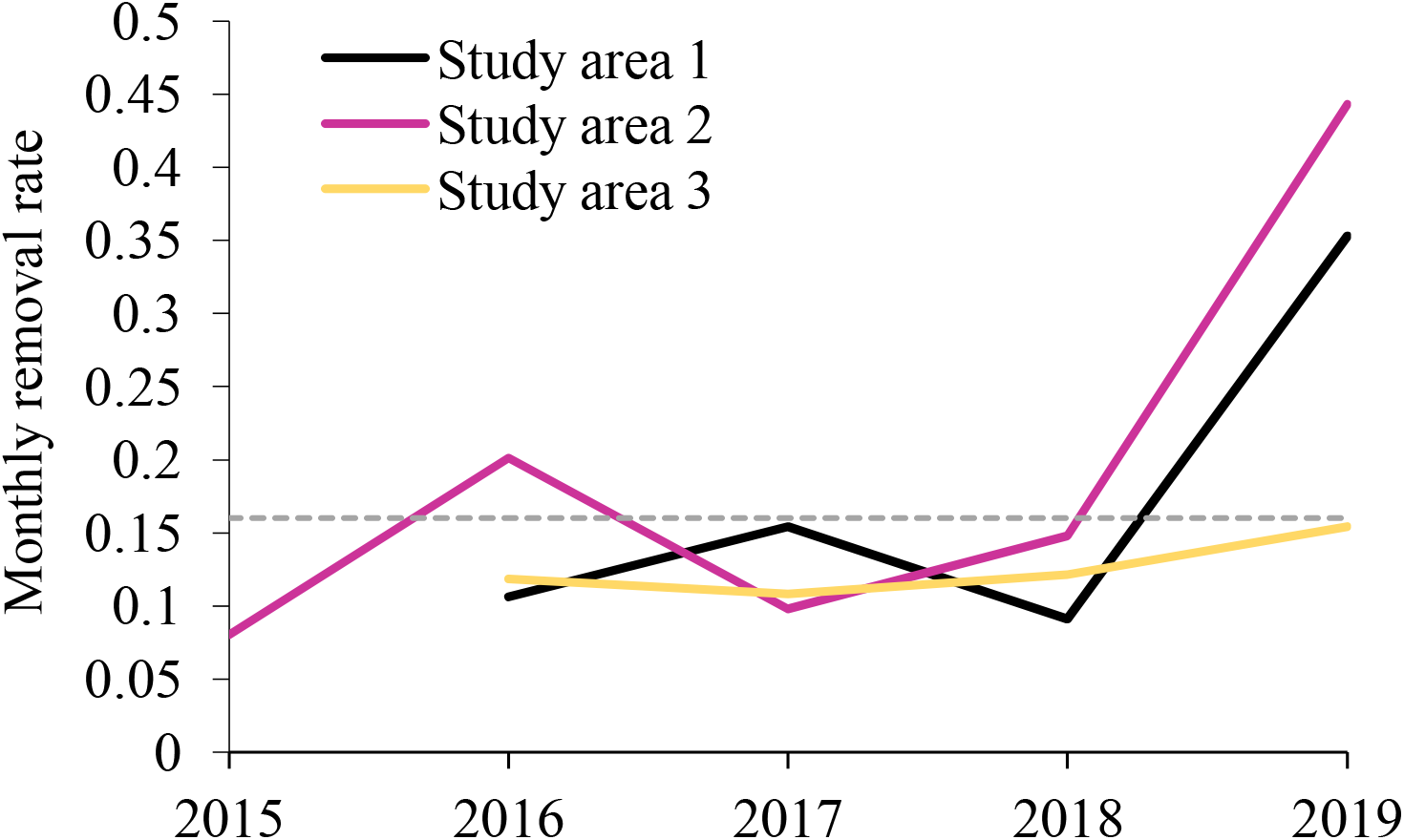
Average proportion of the population removed by month shown by study area and across years. The dashed grey line represents the proportion of the population needed to be removed by month to counteract the population growth. Values above the dashed grey line would result in population declines and values below the dashed grey line would result in population growth.

The capture rates represent the proportion of the population removed for one unit of effort (effort depends on the method, 1 hour in a helicopter, 1 night for trapping) within the area impacted by the removal method. The estimated capture rates for: one trap within a 15 km^2^ area was 0.43 (95% CI: 0.37, 0.51), one hour of aerial gunning in a 40 km^2^ area was 0.18 (95% CI: 0.12, 0.31), and one ground shooting event in a 5 km^2^ area was 0.65 (95% CI: 0.56, 0.77). The entire area covered by all study areas in Mingo NWR was 311.6 km^2^. Therefore, the proportion of the entire population removed (available and captured) by each method was 0.02, 0.023, and 0.010 for trapping, aerial gunning, and ground shooting respectively.

To counteract the population growth, we needed to remove at least 18% of the entire population each month. Based on the estimated average captured rates, this means that for each month either 9 trap nights, 7.2 hours of aerial gunning, or 17 ground shooting events would be needed to maintain a population decrease across all of Mingo NWR. Growth rates did vary in our study over time and as growth rates increase the proportion of the population that must be removed to ensure a population decline will also need to increase.

## DISCUSSION

A primary challenge with invasive species management is balancing the need to monitor change in abundance with the fundamental goal of reducing or eliminating invasive species. Invasive species management incorporates a variety of management removal actions and may be conducted over several months or years, if not perpetually. Currently, no method exists that monitors abundance while accommodating the diversity and intricacies of data from management of invasive animals. We developed a method to estimate changes in abundance over time using only data collected by management removal actions. The method allows for multiple removal methods, occurring non-systematically over a large spatial area and over time. Each monthly abundance estimate is the abundance prior to removal events. By subtraction, we can get the remaining abundance at the end of the removals and calculate the proportion of the population removed at each time point (Davis et al. 2019), thus providing a method for evaluating the impact of management actions on populations over time.

By incorporating multiple removal methods (e.g., aerial gunning, trapping, ground shooting) in our framework, we increase the applicability of removal models to more diverse types of removal data. Management of invasive species involves continually working to improve population suppression methods and therefore, often results in a suite of management approaches being used concurrently to combat a single invasive species (see examples in Clout and Williams 2009, Pepin et al. 2019). Methods that incorporate multiple types of monitoring data not only give more power to estimate ecological states of interest, but also allow managers to learn about the relative value of different data streams (Davis et al. 2019a). In addition to broadening the utility of removal models, the incorporation of multiple removal methods can be used to compare the efficacy of different methods (Bailey et al. 2004, O’Connell Jr et al. 2006). While the intent of our analysis was to use multiple removal approaches to estimate abundance, we can also use these analyses to compare their efficacy. Aerial gunning was the most efficient method at removing feral swine in terms of time, for example, an hour in a helicopter was similar to 1.14 trap nights or 2.24 ground shooting events. However, the costs associated with conducting aerial gunning are considerably higher than running a trap for a night or conducting two ground shooting events and therefore, incorporating costs into these analyses will give a better overall understanding of the cost-effectiveness of different methods. It is also important to consider that the efficacy of different removal methods will vary by habitat, accessibility, time of year, and personnel experience (Davis et al. 2019b). Therefore, the cost-effectiveness of different methods may depend on these factors. Additionally, in this analysis we assumed capture rates were constant within method, but using covariate data from local conditions to inform capture rate parameters would allow us to examine how capture rates within methods vary potentially by time, by habitat, or by personnel (Davis et al. 2019b).

Animals may not be available for detection because they are not behaviorally able to be detected by the detection method (e.g., female birds that don’t sing during breeding bird auditory surveys; Diefenbach et al. 2007) or they may be temporarily unavailable spatially (e.g., submerged whales are unable to be detected by breach surveys even when they are in the study area; Givens et al. 2016). We incorporated availability by considering that an animal may be in the study area but may not be available to a given removal method based on the spatial coverage of that method. The area over which a removal method searches or attracts animals (‘area of influence’ of a method) depends on the method. We used the location information associated with the removal method (e.g., flight track logs, latitude/longitude of trap locations), number of removal events, and information on trap area of influence to estimate the availability by method. We set the area of influence as constant based on previous studies (Davis et al. 2017, McRae et al. 2020), but variation in the area of influence could be included to account for circumstances (e.g., duration of baiting, time of year; Snow and VerCauteren 2019, McRae et al. 2020).

In addition to incorporating multiple removal methods, our approach allows for abundance to vary with open population demographics (i.e. births, deaths, immigration, and emigration), and for population estimation at discrete periods across time, in line with how invasive species management is often conducted. This advance, similar to Link et al. (2018) and Stevens et al. (2020), lets us match the monitoring method with the management method, and continually track population changes in time. The standard removal model would need to be run separately for different removal periods that do not have demographic closure. For a single removal study, it is possible to estimate the abundance pre- and post-removal events and calculate the proportion of the population removed. This is critical information for the monitoring and evaluation of removal activities. By tracking removal events and populations across time, we can monitor populations across time, identify the efficacy of removal efforts, and determine if the ultimate goals of population reduction are being achieved by estimating what the population growth rates are over time.

In our case study we found that populations of feral swine in Mingo NWR had growth rates that varied over the study period but on average intrinsic monthly growth rates (λ) were 1.17. This growth was related to a combination of births and immigration. This growth rate is higher than expected based on estimates of annual growth rate of feral swine from births alone (Timmons et al. 2012, Mellish et al. 2014). However, these are local-level estimates of population growth that may include immigration within the overall refuge and surrounding areas in response removal efforts. Depopulating an area can potentially cause a vacuum effect where animals will immigrate into an area where animals were recently removed especially if that area contains desirable habitat (Killian et al. 2007). Removal efforts need to be able to remove more animals than those produced by births and immigration to ensure the population is going to decline. Based on the estimates of monthly population growth, approximately 18% of the population in Mingo NWR should be removed monthly to counteract population growth and see a population decline over time. However, as these rates of population growth change over time the effort needed to control the population will change as well. Therefore, continued monitoring using this approach is recommended. If these high levels of growth are due in part to the vacuum effect, then immigration rates may decline after initial pulses of immigration and thus the growth rate within the refuge will decrease. This reflects what we observed in Mingo NWR, as there was a sharp increase in growth in response in intensive removal efforts but with sustained removal efforts the growth rates fell considerably near the end of the study.

This dynamic removal model that incorporates multiple removal methods is more broadly applicable to management conditions than static or single-method approaches. However, management data are not designed for population estimation and therefore often do not meet the assumptions necessary to apply abundance estimation methods. Model requirements include population closure during sampling (here 1 month), all animals are equally catchable, capture rates are constant with respect to effort, and that sampling periods are short relative to the open intervals (Zippin 1958, Pollock 1982). As management removals are conducted based on personnel and resource availability and often not according to a systematic design, we specified a sampling period as the duration from the first removal event to the last removal event within a given month, and the open period was the time from the last removal event in one month to the first removal event in the next. This means that the open period may range from 1 day to over 30 days. We adjusted the growth rate calculations to adapt to this variable time period. However, the periods of closure may be violated if they are closer to a month in length than a week. Kendall (1999) found that violations of closure did not bias estimates if movement on and off the study area is random for capture-recapture abundance estimates. He also found that if movement was only on or only off of the study area the abundance estimates were unbiased relative to the superpopulation (Kendall 1999). Kendall (1999) did find that bias was introduced if movement was Markovian (based on the status the previous time period). The impacts on removal estimates may be different from capture-recapture estimates, but in general violations of closure assumptions lead to overestimation of abundances. Although we would like to be accurate in our abundance estimation, for invasive species it can be more beneficial to overestimate populations as underestimation could lead to a decline in management effort which may allow the population to rebound.

In acknowledgement that management data are not designed for population estimation and therefore often do not meet the assumptions necessary to apply abundance estimation methods, we tested the robustness of this removal approach to violations of these assumptions with extensive simulations. Even in situations when the population estimates were biased high or low, the model still captured the general pattern of population change well, suggesting that trends in population can be monitored using this approach. Auxiliary studies may be needed if the conditions for unbiased estimates outlined above are not met. These simulations help provide guidelines on when this dynamic removal approach would be appropriate to use on management data. Additionally, removal models are known to perform poorly when the number of animals removed per pass is small either due to low abundance or capture probabilities (Seber and Whale 1970, Davis et al. 2016). Therefore, this method is better applied to populations in reduction phases and not for populations nearing elimination. This approach can be used in an adaptive management framework that allows the monitoring data to inform when abundance estimation should be used and when analyses should switch to occupancy modeling (i.e., estimating the probability of presence rather than abundance).

Objectives associated with invasive species management may range from elimination of the invasive species, to maintaining a low acceptable level of the invasive species, or to focus on damage reduction of the invasive species. For any of these objectives some form of monitoring needs to be conducted to evaluate if the resources being spent are achieving the desired objective. By using the approach we developed, managers can evaluate their control actions without diverting resources from additional control activities and show evidence of the impact of their actions for stakeholders. Additionally, our approach can be used to evaluate the effectiveness of management actions and be combined with economic approaches to determine the cost effectiveness of different management actions and optimal control strategies (Pepin et al. 2020).

## ACKNOWLEDGEMENTS

We would like to thank Wildlife Service and National Parks Service personnel. The research was funded by the APHIS National Feral Swine Damage Management Program. Any use of trade, firm, or product names is for descriptive purposes only and does not imply endorsement by the U.S. government. This research did not receive any specific grant from funding agencies in the public, commercial, or not-for-profit sectors.

